# A novel auxin-inducible degron system for rapid, cell cycle-specific targeted proteolysis

**DOI:** 10.1101/2021.04.23.441203

**Authors:** Marina Capece, Anna Tessari, Joseph Mills, Gian Luca Rampioni Vinciguerra, Chenyu Lin, Bryan K McElwain, Wayne O. Miles, Vincenzo Coppola, Dario Palmieri, Carlo M. Croce

**Affiliations:** Department of Cancer Biology and Genetics, The Ohio State University, 43210, Columbus, OH, USA; The Ohio State University Wexner Medical Center and Comprehensive Cancer Center, 43210, Columbus, OH, USA; Gene Editing Shared Resource, The Ohio State University Wexner Medical Center and Comprehensive Cancer Center, 43210, Columbus, OH, USA

**Keywords:** Auxin-inducible degron, OsTIR1, Cell Cycle

## Abstract

The OsTIR1/auxin-inducible degron (AID) system allows “on demand” selective and reversible protein degradation upon exposure to the phytohormone auxin. In the current format, this technology does not allow to study the effect of acute protein depletion selectively in one phase of the cell cycle, as auxin similarly affects all the treated cells irrespectively of their proliferation status. Therefore, the AID system requires coupling with cell synchronization techniques, which can alter the basal biological status of the studied cell population. Here, we introduce a new AID system to Regulate OsTIR1 Levels based on the Cell Cycle Status (ROLECCS system), which induces proteolysis of both exogenously transfected and endogenous gene-edited targets in specific phases of the cell cycle. This new tool paves the way to studying the differential roles that target proteins may have in specific phases of the cell cycle.

## Introduction

The cell-division cycle, also known as cell cycle, is the fundamental, precise and complex process at the basis of life and physiological processes such as development, tissue growth, homeostasis, regeneration, and aging in multicellular organisms^1, 2^. In mitotic cells, the division into two daughter cells (cytokinesis) occurs after the parental cell undergoes the semiconservative synthesis (S phase) of a new copy of its entire genome, followed by the mitotic chromosomal segregation (M phase). Two gap phases, G0/G1 and G2, precede the S and M phases, respectively^1–3^.

The mechanisms leading to and controlling DNA replication and segregation are historically among the most studied and understood processes happening throughout cell division^4^. Critical molecular players involved in cell cycle regulation and control have been identified based on the effect that their mutation, overexpression, or silencing have on genome replication, either in physiological conditions or in response to DNA damaging agents^3, 5, 6^.

However, genomic DNA is not the only cellular component undergoing dramatic changes during cell cycle progression. Proteins, organelles, and cellular membranes experience profound modifications to allow the appropriate segregation of all the required materials in the daughter cells^7, 8^. One obvious paradigm is constituted by the nuclear membrane, which disassembles immediately before cells enter mitosis to be promptly re-assembled at the completion of the cell division cycle^7–9^. During this process, the nuclear content and proteins of the nuclear pore complexes are released in the open cytoplasm, and novel protein-protein interactions can take place^10–12^. Hence, it could be assumed that virtually any cellular protein might become part of alternative multi-protein complexes and perform different biological tasks, such as preserving DNA integrity^13^ or cytoskeletal dynamics^14^.

To date, existing technical limitations have prevented an appropriate discrimination of phase-specific protein functions, especially in physiological conditions. Insights into the cell cycle regulatory networks were initially obtained by analyzing cells synchronized in specific phases of the cell cycle. However synchronization is routinely achieved by exposing cells to stress conditions, such as serum starvation, inhibition of DNA synthesis, or by disrupting microtubule dynamics^15, 16^.

A major advancement in the field was represented by the development of the FUCCI (Fluorescent Ubiquitination-based Cell Cycle Indicator) system^17–19^. This technology is based on the enzymatic activity of the two E3 ubiquitin ligases, APC^Cdh1^ and SCFSkp2 ^19–22^, involved in the control and proteasomal degradation of the replication origins licensing factors Geminin (targeted by APC^Cdh1^) and Cdt1 (targeted by SCF^Skp2^)^23^. By fusing Cdt1 and Geminin with a variety of fluorescent proteins, a number of tools were engineered to accurately discriminate the cell cycle status of individual cells, either microscopically or by flow cytometry^19, 24–27^, both *in vitro* and *in vivo*.

This novel approach provides an efficient way to identify, visualize and select cells in specific phases of the cell cycle. However, it must be still combined with other technologies to perturb the levels of the protein of interest (POI) to assess its biological role throughout the cell cycle. Despite the advancements of genetic tools such as CRISPR/Cas9-based gene editing^28^, gene silencing^29^, or inducible gene expression approaches^30^, none of these systems displays readiness of activity compatible with the kinetics of cell cycle progression. Conversely, an alternative to obtain rapid degradation of the POI is represented by targeted proteolysis using PROteolysis-Targeting Chimeras (PROTACs), or polypeptide tags (also known as degrons)^31, 32^. In one of the most commonly used degron systems, the POI is fused with an Auxin-Inducible Degron (AID) sequence, such as the 7 kDa degron termed mini-AID (mAID), in cell lines expressing the *Oryza sativa* TIR1 (*Os*TIR1) F-box protein^33, 34^. When the phytohormone auxin is provided, OsTIR1 binds the mAID-POI and induces its quick proteasomal degradation^33, 34^. However, despite their speed, reversibility, and fine-tuning, degron-based systems still lack cell cycle phase-specificity and require conventional cell synchronization^35^.

Here, we report the engineering the “Regulated OsTIR1 Levels of Expression based on the Cell-Cycle Status” (ROLECCS) technology that combines the AID and the FUCCI systems. In this new tool, the *Os*TIR1 protein is fused to the fluorescent indicator mEmerald and the FUCCI tags Cdt1/Geminin, which are responsible for the restricted G1 and S/G2 expression, respectively. Upon auxin treatment, only the cells expressing the fusion-protein OsTIR1-mEmerald-Cdt1/Geminin, (i.e. in the desired cell cycle phase) degrade the mAID-POI.

We developed this technology to provide a unique tool for studying the differential role of POIs throughout cell cycle progression, taking advantage of its cell cycle phase-specificity, rapidity, reversibility, and low overall perturbation of other biological processes.

## Results

### Designing and engineering a cell cycle phase-specific OsTIR1

To engineer a cell cycle phase-specific degron system, we generated a variant of the mAID system where the expression of OsTIR1, necessary for the recognition and degradation of the mAID-tagged protein upon auxin exposure, was dependent on the G0/G1 or S/G2/M phase.

In our design, the OsTIR1 coding gene was fused in-frame with a mEmerald fluorescent reporter (a brightly fluorescent monomeric variant of GFP)^36^ that allows the identification of cells expressing these constructs. Then, we added the sequences corresponding to either human Cdt1 (aa 30-120) or Geminin (aa 1-110) to restrict OsTIR1 expression to different phases of the cell cycle, like in the FUCCI system. For convenience, the hCdt1 (30-120) and hGeminin (1-110) tags are indicated hereafter as Cdt1 and GEM, respectively. We also generated a construct where no additional tag was added, to allow OsTIR1-mEmerald expression independently on the phase of the cell cycle (**Figure 1**).

**Figure 1.**
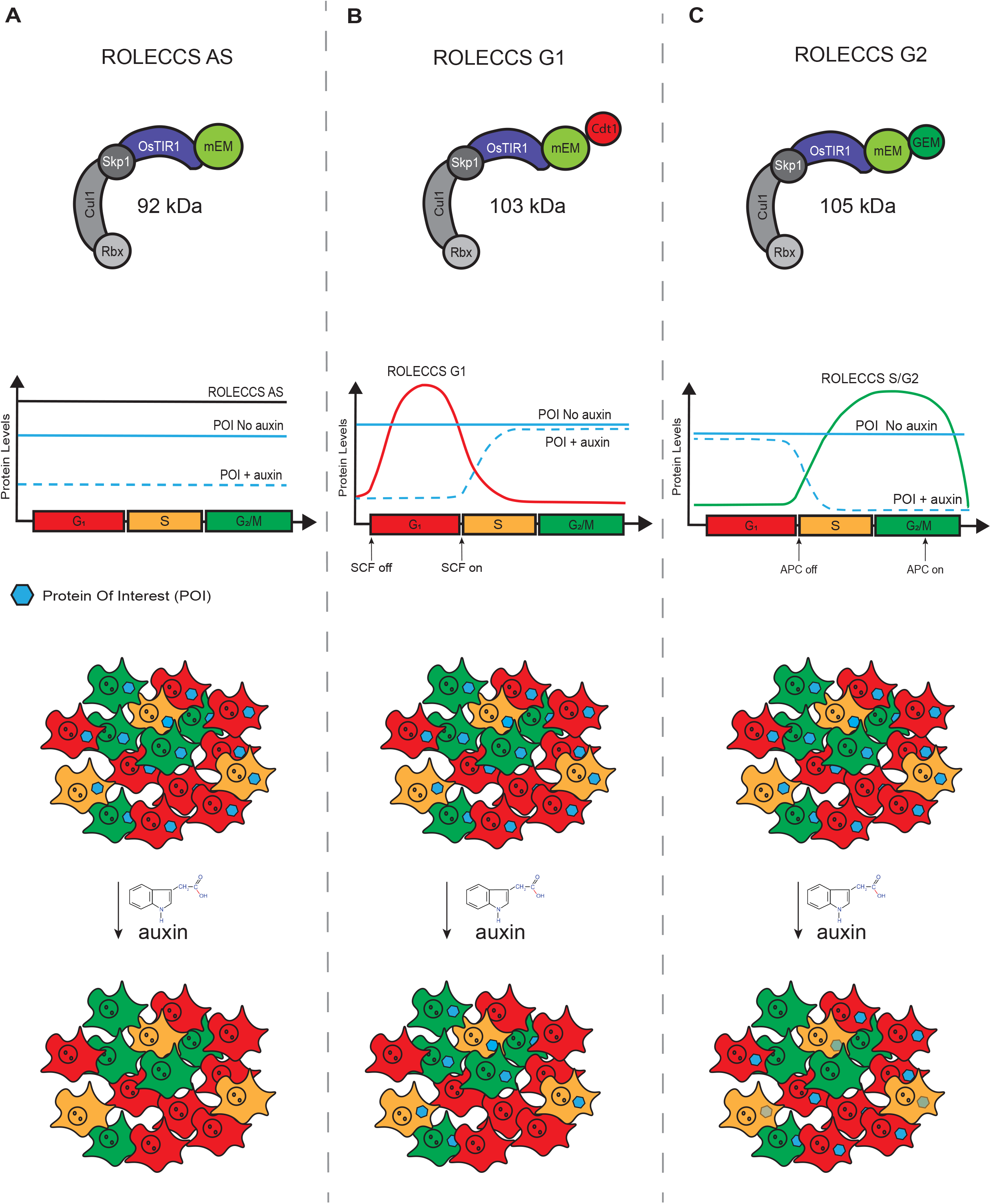
Schematic representation of the design for Regulated OsTIR1 Levels of Expression based on the Cell Cycle Status (ROLECCS) variants. (A) OsTIR1-mEmerald protein (asynchronous ROLECCS, ROLECCS AS, 92 KDa) is stably expressed throughout the cell cycle. Upon auxin treatment, OsTIR1 enzymatic activity elicits the degradation of the mAID-tagged protein of interest (POI) in any cell, independently of the cell cycle status. (B) The expression of the ROLECCS G1 variant (OsTIR1-mEmerald-Cdt1, 103 KDa) is restricted to the G1/early S phase by the presence of the Cdt1 tag, when the SCF^Skp^^2^ E3 ligase activity is off. This, in turn, leads to auxin-dependent ubiquitylation and proteasome degradation of mAID-tagged POIs. In cells transitioning during S, G2 and M phases, SCF^Skp^^2^ activity is naturally restored, leading to ROLECCS G1 degradation by ubiquitylation, and stabilization of the POI even in the presence of auxin. (C) The Geminin tag of the ROLECCS G2 variant (OsTIR1-mEmerald-GEM, 105 KDa) ensures its restricted expression during the late S-G2-M phase, as APC^Cdh^^1^-mediated ubiquitylation and degradation is rapidly triggered during M/G1 transition. Therefore, auxin treatment induces degradation of the POI exclusively in cells going through the late S-G2-M phase of the cell cycle.

In our design, engineered variants of OsTIR1-mEmerald, OsTIR1-mEmerald-Cdt1, and OsTIR1- mEmerald-GEM genes are actively transcribed throughout the cell cycle. However, the presence of the Cdt1 and the Geminin tags determine the Regulated OsTIR1 Levels of Expression based on the Cell Cycle Status (ROLECCS system). We predicted that the OsTIR1-mEmerald protein would be stably present throughout the cell cycle. Therefore, auxin treatment would trigger OsTIR1 enzymatic activity and degradation of the mAID-tagged protein of interest in any cell, independent of the cell cycle status (from now on: asynchronous ROLECCS, ROLECCS AS) (**Figure 1A**).

On the other hand, the presence of OsTIR1-mEmerald-Cdt1 (from now on: ROLECCS G1) protein would be restricted the G1/early S phase, because ubiquitylation by SCF^Skp^^2^ E3 ligase leads to its prompt degradation during S-phase transition. Thus, addition of auxin would lead to OsTIR1- mediated proteasomal degradation of the POI exclusively in those cells in G1/S phase during the treatment (**Figure 1B**).

Similarly, presence of OsTIR1-mEmerald-GEM (from now on: ROLECCS G2) protein would be restricted during the late S-G2-M phase, peaking during the G2, as APC^Cdh^^1^-mediated ubiquitylation and degradation is rapidly triggered during M/G1 transition. Consequently, auxin treatment would cause degradation of the POI exclusively in cells going through the late S-G2-M phase of the cell cycle during the treatment (**Figure 1C**). To provide flexibility to the system and make it usable in different paradigms, the three CMV-driven ROLECCS constructs were subcloned in *ad hoc* vector^33^ that allows either transient expression or CRISPR/Cas9-mediated integration into the *AAVS1* safe harbor site of the human genome (**Supplementary Figure 1A-C**).

### ROLECCS G1 and ROLECCS G2 expression during cell cycle

To demonstrate that ROLECCS G1 and ROLECCS G2 expression is restricted to specific phases of the cell cycle, we first assessed their relative abundance in transiently transfected HEK-293 cells. As shown in **Figure 2A**, each ROLECCS construct was abundantly expressed at 72h from transfection. Unlike the previously published FUCCI probes^19^, all the ROLECCS proteins were present both in the nucleus and in the cytoplasm of transfected cells.

**Figure 2.**
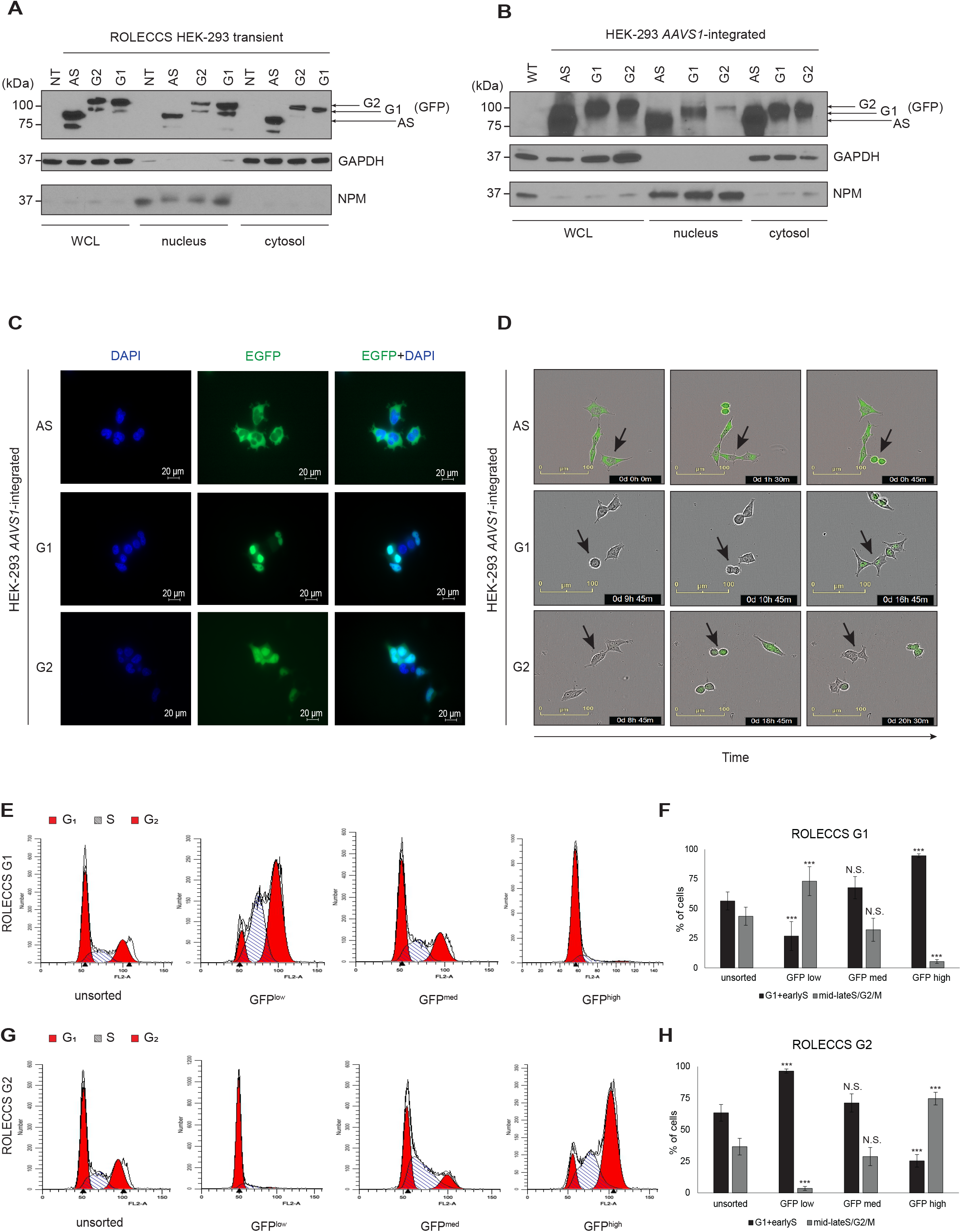
Characterization of ROLECCS AS, G1, and G2 cellular distribution and during cell-cycle progression. (A-B) Representative WB analysis of nuclear/cytoplasmic distribution of ROLECCS proteins upon transient (72hrs) transfection (A) or *AAVS1* integration (B) in HEK-293 cells. ROLECCS AS, ROLECCS G1, ROLECCS G2 (see main text) were detected using anti-GPF antibody, that recognizes the mEmerald tag of the proteins (see arrows). Nucleophosmin (NPM) and GAPDH antibodies were used as loading and purity control for nuclear and cytoplasmic soluble protein fractions, respectively. Not transfected (NT) or wild-type (WT) HEK-293 were used as negative control. WCL indicates Whole Cell Lysate. (C) Direct fluorescence images of HEK-293 *AAVS1*-integrated clones. DAPI staining (blue) was used to label nuclei, EFGP (green) signal was detected from ROLECCS variants (AS, G1, G2). (D) Time-frame pictures of duplicating HEK-293 *AAVS1*-integrated clones. Note the cell cycle-dependent changes in fluorescence of specific ROLECCS variants (AS, G1, G2) (green). Arrows indicate cells that are completing a cell cycle. (E, G) Cell-cycle distribution histograms of HEK-293 *AAVS1*-integrated clones expressing ROLECCS G1 and G2, obtained by propidium iodide staining and flow cytometry analysis. Red peaks indicate G1 and G2 phase, stripes indicate S phase. Cells were prior sorted based on GFP levels (GFP^low^, GFP^med^, GFP^high^), as described in Supplementary Figure 2. Not sorted (unsorted) populations are reported for comparison. Data are representative of four independent experiments. (F-H) Quantification of experiments reported in E and G. GFP^low^, GFP^med^, GFP^high^ subpopulations were analyzed for cells composition as percentage of cells in G1+earlyS and cells in mid-lateS/G2/M, using ModFit software v5.0. Error bars indicate mean ± SD. *** p < 0.001, N.S. not significant. Statistics (two-tailed t-test) is calculated *versus* respective unsorted populations. Data are the average of four independent experiments.

To obtain cell populations expressing a uniform level of the ROLECCS proteins, we generated stable HEK-293 cell lines. To this aim, the AAVS1 ROLECCS vectors were integrated into the *AAVS1* safe harbor locus by CRISPR/Cas9-mediated gene knock-in. Also in this case, sustained and ubiquitous expression of the ROLECCS proteins was observed by nuclear/cytoplasmic protein fractionation (**Figure 2B**), and direct fluorescence imaging (**Figure 2C**).

Next, we aimed to demonstrate that ROLECCS G1 and ROLECCS G2 protein levels oscillate reciprocally during cell cycle transition, as expected. Live cell imaging was performed on ROLECCS AS, G1 and G2 knock-in HEK-293 to monitor the green fluorescence in real time. **Figure 2D** (top) and **Supplementary Video 1** show that ROLECCS AS expression did not change during a full cell cycle. Conversely ROLECCS G1 was not visible in actively dividing cells (**Figure 2D, middle and Supplementary Video 2**), becoming detectable immediately upon completion of cell division. Finally, ROLECCS G2 was visible only in actively dividing cells, with the fluorescence intensity peaking at G2/M transition (**Figure 2D, bottom and Supplementary Video 3**).

To orthogonally validate ROLECCS G1 and ROLECCS G2 as cell cycle indicators, we sorted *AAVS1*-integrated ROLECCS HEK-293 based on their green fluorescence level and cellular complexity (Side Scatter, SSC) (**Supplementary Figure 2A-D**), as described in the Methods section. DNA content analysis demonstrated that GFP^high^-sorted ROLECCS G1 population mostly comprised cells in the G1/early S phase (94.6±1.6%), in comparison with GFP^med^ and GFP^low^ sorted populations (67.7±9.4% and 26.9±12.2% respectively)(**Figure 2E and 2F**). Conversely, GFP^high^ ROLECCS G2 population showed a significant enrichment in late S/G2 phase cells (74.64±5%), compared to GFP^med^ and GFP^low^ sorted cells (28.8±7.2% and 3.5±1.5%, respectively) (**Figure 2G and 2H**). Conversely, unsorted ROLECCS G1 and ROLECCS G2 populations displayed cell cycle distribution typical of unsynchronized HEK-293 cells.

Altogether, our findings indicate that engineered ROLECSS G1 and G2 protein levels are efficiently restricted to specific phases of the cell cycle, as expected.

### Biological activity of ROLECCS

The addition of large tags to proteins might affect their biological activity^37^. Therefore, we wanted to assess that the enzymatic activity of OsTIR1-containing SCF complexes was not hampered by the mEmerald-Cdt1 and mEmerald-GEM tags of the ROLECCS G1 and G2, respectively. We transiently transfected *AAVS1*-integrated ROLECCS AS, ROLECCS G1, and ROLECCS G2 HEK-293 cells with a mAID-mCherry fluorescent reporter and measured its protein levels. As shown in **Supplementary Figure 3**, mAID-mCherry levels were appreciably reduced upon auxin treatment when performed at 8h after reporter vector transfection, indicating that the biological activity of OsTIR1 was preserved. However, transient transfection could lead to multiple sub- populations of ROLECCS-expressing cells with different levels of mAID-mCherry due to inconsistent transduction. Moreover, at later time points, auxin-dependent degradation of mAID-mCherry was negligible (not shown). Therefore, we hypothesized that an overexpressed target could be efficiently degraded only if the molar ratio between the POI and the ROLECCS was favorable to the latter, as in the very first hours (<8h) after transfection. Accordingly, other groups have generated All-in-One systems to achieve equimolar levels of OsTIR1 and its targets^38^.

As shown in **Figure 3A-C**, we created 3 all-in-one lentiviral vectors (pLentiROLECCS AS, G1, and G2) in which the ROLECCS proteins were fused with mAID-mCherry using an autoproteolytic P2A sequence^39^. This approach allows the simultaneous and equimolar expression of both OsTIR1 and its target which, upon translation, are released as independent molecules. **Figure 3D** shows that transient and stable transfection of pLentiROLECCS vectors led to sustained expression of both the ROLECCS proteins and of mAID-mCherry.

**Figure 3.**
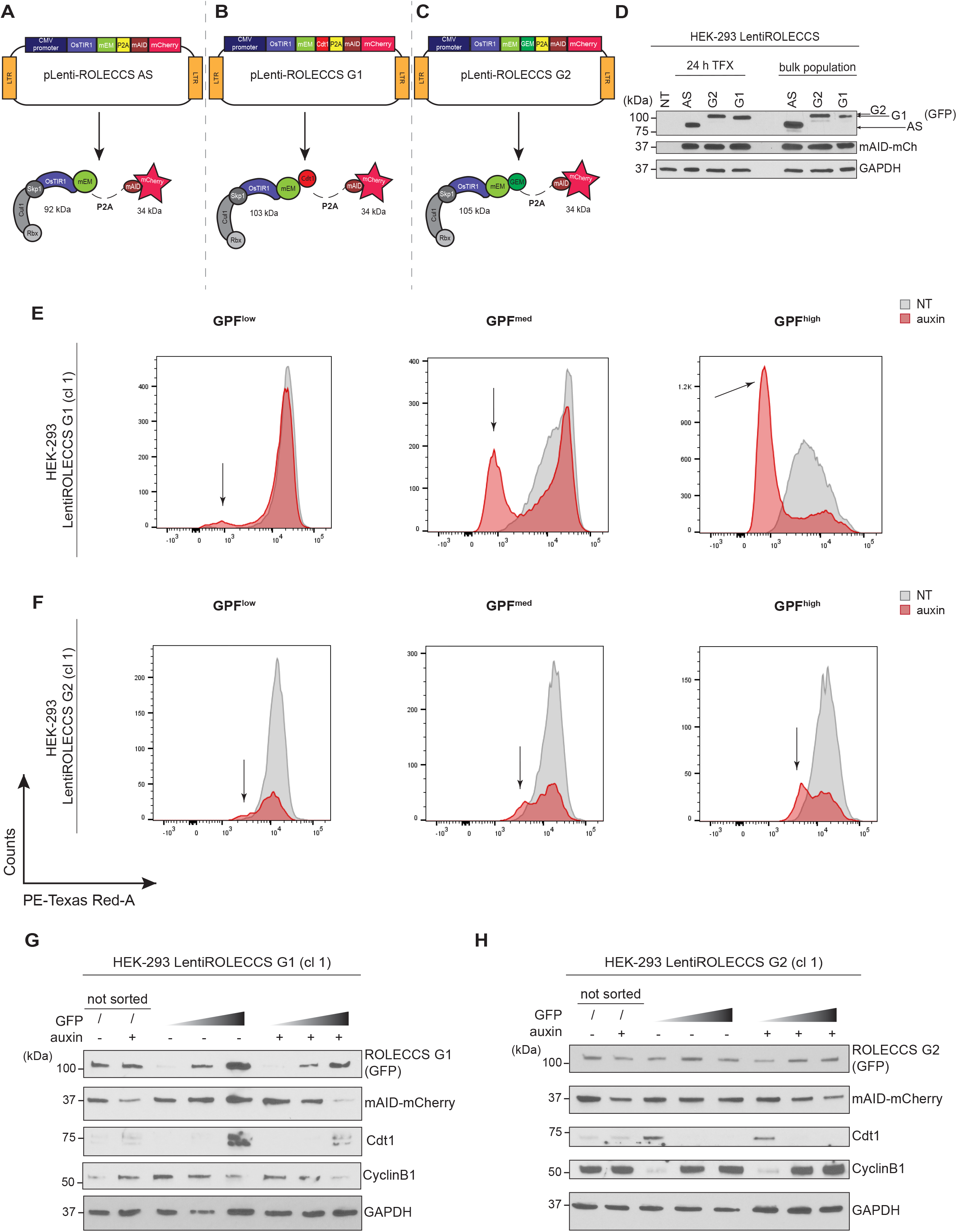
Biological activity of ROLECCS proteins. (A-C) Schematic representation of lentiviral vectors (pLentiROLECCS AS, G1, and G2) and their corresponding translated proteins with respective molecular weight. (D) WB analysis of transient (24 hours) and stable transfection (bulk population) of pLentiROLECCS vectors in HEK-293 cells. Anti-GPF antibody was used to detect ROLECCS proteins (see arrows), anti-mCherry antibody was used to detect mAID- mCherry. Not transfected HEK-293 (NT) and GAPDH were used as negative and loading control respectively. (E-F) Representative histograms of red fluorescent intensity (PE-Texas Red-A *versus* counts) of GFP sorted subpopulations for LentiROLECCS G1 (E) and LentiROLECCS G2 (F) HEK-293 clones. Arrows indicate cells shifting toward lower intensity of red fluorescence upon auxin treatment. Data are representative of four independent experiments. (G) WB analysis of HEK-293 cells transfected with pLentiROLECCS G1, (clone 1, cl 1) after sorting. Cells were treated with auxin or left untreated for one hour and then sorted based on increasing GFP intensity (ascending grey gradient triangle). Membrane was probed with anti-GFP antibody for ROLECCS G1 detection and mCherry antibody for mAID-mCherry detection. Cdt1 and CyclinB1 were used as G1 phase and G2 phase specific markers, respectively. GAPDH was used as loading control. Not sorted (unsorted) cells were loaded for comparison. (H) WB analysis of HEK-293 cells transfected with pLentiROLECCS G2 (clone 1, cl 1), after sorting. Treatments, sorting and antibodies were performed as in G. Blots are representative of at least two independent experiments.

Next, we treated HEK-293 cells stably expressing LentiROLECCS G1 or LentiROLECCS G2 with auxin for 1h. Cells were sorted based on their GFP fluorescence intensity and SSC. Red fluorescent intensity (from mAID-mCherry) was simultaneously quantified on GFP^low^, GFP^med^, and GFP^high^ populations. **Figure 3E-F** show that downregulation of the mCherry fluorescence was specifically achieved in GFP^med^ and GFP^high^ populations upon auxin treatment.

Western blot analysis confirmed that GFP^med^ and GFP^High^ sorted cell populations expressed the highest levels of ROLECCS G1 and ROLECCS G2 (**Figure 3G-H**). Notably, highest levels of ROLECCS G1 corresponded to highest expression of Cdt1 (a G1-specific marker, frequently identified as a doublet corresponding to Cdt1/phosphoCdt1^40^) and to the lowest levels of Cyclin B1 (a late-S/G2 marker). Importantly, downregulation of the target mAID-mCherry was only observed in sorted GFP^high^ ROLECCS G1 cells upon auxin treatment, and not in the untreated or GFP^low^ auxin-treated controls (**Figure 3G)**. Similarly, ROLECCS G2 accumulation was observed in cell populations displaying highest levels of Cyclin B1 and lowest levels of Cdt1, but target downregulation was only observed upon auxin treatment (**Figure 3H**). These results were confirmed using 2 independent HEK-293 LentiROLECCS clones (**Supplementary Figure 4A-B**).

Taken together, our data indicate that the ROLECCS system allows temporally-restricted selective degradation of mAID-tagged targets based on the cell cycle phase.

### The ROLECCS system allows cell cycle phase-specific downregulation of endogenous proteins

It has been previously demonstrated that the mAID system is suitable for the downregulation of endogenous protein, when the gene of interest is modified by CRISPR/Cas9-mediated knock-in to include the mAID sequence^33^. We aimed to demonstrate that the ROLECCS system allows the same capability, but only in the phase of the cell cycle of interest. We decided to test whether the ROLECCS system could accomplish the cell cycle phase-specific downregulation of TP53, a well-known transcriptional factor playing a central role in the control of cell cycle progression, especially in response to DNA-damaging agents^41, 42^. Moreover, one of the main mechanisms of physiological negative regulation of TP53 is its MDM2-mediated ubiquitylation and proteasomal degradation^41^. Therefore we postulated that ROLECCS-mediated synthetic degradation of TP53 could represent a valid alternative to its physiological mechanism of regulation.

First, we generated HCT116 cell lines where both wild-type *TP53* alleles were modified by CRISPR/Cas9-mediated knock-in (HCT116 TP53-mAID-mCherry). For gene editing purposes, the stop codon of the endogenous *TP53* gene was replaced by a mAID-mCherry fusion cassette (as described in Methods section) (**Figure 4A**, **Supplementary Figure 5A**). Appropriate editing by site-specific integration of the donor cassette was verified by PCR using integration-specific primer sets, as shown in **Supplementary Figure 5B-C**.

**Figure 4.**
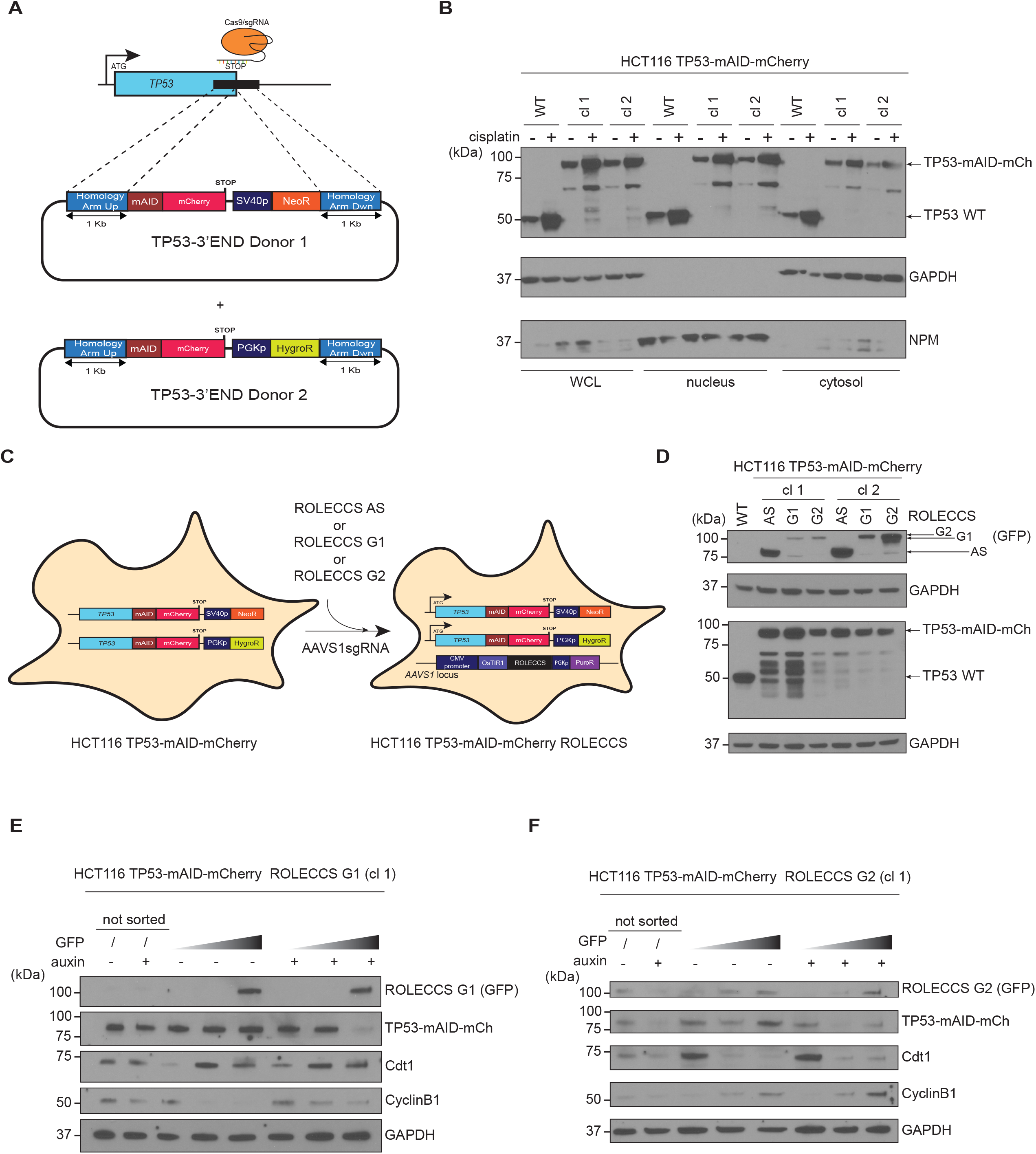
ROLECCS system downregulates endogenous proteins in a cell cycle-specific fashion. (A) Diagram of *TP53* gene editing strategy in HCT116 via CRISPR/Cas9-mediated knock-in. The stop codon was replaced by mAID-mCherry fusion cassette, cloned between 1-kb long Homology Arms. To achieve targeting of both *TP53* alleles, two donor plasmids (TP53- 3’END Donor 1 and Donor 2) were used, bearing Neomycin (NeoR) or Hygromycin (HygroR) resistance genes, respectively. The antibiotic resistance genes are under the transcriptional control of independent promoters (SV40 and PGK, respectively). (B) WB analysis of nuclear/cytoplasmic distribution of TP53 protein (TP53-mAID-mCherry, 87 kDa) in HCT116 TP53-mAID-mCherry clone 1 (cl 1) and clone 2 (cl 2). HCT116 wild type (WT) were loaded as control for TP53 activation upon cisplatin (20 µM) treatment for 48 hours. NPM and GAPDH were used as purity and loading controls for nuclear and cytoplasmic soluble protein fractions, respectively. WCL indicates Whole Cell Lysate. Images are representative of two independent experiments. (C) Schematic illustration of the generation of HCT116 TP53-mAID-mCherry ROLECCS cells by introducing the transgenes ROLECCS (AS/G1/G2) at the safe-harbor *AAVS1* locus via CRISPR/Cas9 in HCT116 TP53-mAID-mCherry. (D) WB analysis of characterization of HCT116 TP53-mAID-mCherry with *AAVS1*-integrated ROLECCS variants (AS/G1/G2). ROLECCS AS, ROLECCS G1, and ROLECCS G2 were detected using anti-GPF antibody (see arrows), TP53 wild type (WT) and TP53-mAID-mCherry (TP53-mAID-mCh) were detected using anti-TP53 antibody. GAPDH was used as loading control. HCT116 wild type (WT) were loaded for comparison. Two independent HCT116 TP53-mAID-mCherry ROLECCS clones (cl 1, cl 2) were analyzed. Images are representative of two independent experiments. (E) WB analysis of HCT116 TP53-mAID-mCherry *AAVS1*-edited with ROLECCS G1, clone 1 (cl 1) after sorting. Cells were treated with auxin or left untreated for one hour and then sorted for GFP intensity (ascending grey gradient triangle). Membrane was probed with anti-GFP antibody for ROLECCS G1 detection and mCherry antibody for TP53-mAID-mCherry (TP53-mAID-mCh) detection. Cdt1 and CyclinB1 were used as G1 phase and G2 phase specific markers, respectively. GAPDH was used as loading control. Not sorted (unsorted) cells were loaded for comparison. (F) WB analysis of HCT116 TP53-mAID-mCherry *AAVS1*-edited with ROLECCS G2, clone 1 (cl 1) after sorting. Treatments, sortings, and antibodies are the same as shown in panel E. Blots are representative of two independent experiments.

Western blot analysis showed that gene-edited TP53 had a marked molecular size increase (final predicted molecular weight ∼87KDa, compared to WT TP53, 53KDa), due to the presence of the mAID and mCherry tags. Of note, when probed with a TP53-specific antibody, edited clones displayed additional lower molecular weight bands, possibly due to an unstable linker sequence, previously described at the N-terminal domain of mCherry^43^. Importantly, gene-edited TP53 was still upregulated by DNA damaging agents such as cisplatin treatment, and its nuclear and cytoplasmic localization followed the expected distribution pattern^41, 42^ (**Figure 4B**). The additional lower bands displayed a similar trend upon genotoxic stress.

Next, we further edited HCT116 TP53-mAID-mCherry cells inserting the ROLECCS constructs in the *AAVS1* safe harbor site, generating HCT116 TP53-mAID-mCherry ROLECCS cell lines (**Figure 4C**). As shown in **Figure 4D**, sustained expression of ROLECCS AS, G1, and G2 with the expected molecular weight was achieved in at least 2 independent clones. Since this analysis was performed on asynchronously growing HCT116 TP53-mAID-mCherry ROLECCS cells, the three ROLECCS constructs apparently displayed different expression levels. However, these differences are likely due to the fact that ROLECCS G1 and ROLECCS G2 are expressed only in phase-specific cell subpopulations, while ROLECCS AS is equally expressed throughout the cell cycle. Nonetheless, since we did not perform relative comparisons between ROLECCS AS and ROLECCS G1 or G2, these differences did not affect downstream analyses. We also noticed a mild reduction in the levels of edited TP53 in comparison with parental HCT116 cells, compatible with the partial leakiness observed for the AID system^44, 45^. For this reason, for all the functional studies, cells were pre-treated with auxinole (as described in the Methods section), a previously reported inhibitor of OsTIR1^44^, to neutralize the activity of the ROLECCS system in the absence of auxin.

Finally, to validate that the ROLECCS system could allow cell-cycle specific target degradation of an endogenous target, we treated HCT116 TP53-mAID-mCherry ROLECCS G1 and G2 cells with auxin. At 1h after auxin treatment, cells were sorted (**Supplementary Figure 6A-D**) based on their green fluorescence and SSC, as described in the Methods section. As shown in **Figure 4E-F**, TP53 downregulation was noticeable in unsorted populations both in ROLECCS G1- and G2-expressing cells. However, upon sorting, we observed that TP53 downregulation upon auxin treatment was only achieved in GFP^med^ and GFP^high^ sorted populations, in comparison with GFP^low^ cells for both ROLECCS constructs. Importantly, Cdt1 and Cyclin B1 levels confirmed that ROLECCS G1 GFP^med^ and GFP^high^ represented a cell population enriched in G1/early S phase of the cell cycle. On the other hand, ROLECCS G2 GFP^med^ and GFP^high^ cells were mostly representing cells in the late S/G2 phase. Similar results were obtained using two independent clones for each ROLECCS proteins (**Supplementary Figure 7A-B**).

These data indicate that the ROLECCS system can be used to achieve the phase-specific downregulation of an endogenous target, appropriately gene edited to include a mAID tag.

## Discussion

The temporal discrimination of protein functions is critical to fully understand how the same factor might carry out different tasks during different phases of the cell cycle, ultimately leading to diverse biological outcomes. Therefore, “timing is everything”^46^.

The development of mAID systems has allowed sharp and quick modulation of the levels of a protein of interest^32, 47, 48^. Considering the relatively short duration of each phase of the cell cycle, a rapid depletion of the protein of interest is of paramount importance. This is usually not achievable with traditional methods (e.g. RNA interference, gene knockout) for gene silencing. Moreover, these procedures are not promptly reversible, and often generate a phenotype that can potentially produce confounding results due to the absence of the POI throughout multiple cell cycle phases.

In this report, we introduced a novel tool to rapidly and reversibly regulate levels of virtually any protein in a cell cycle phase-specific manner, using the “Regulated *Os*TIR1 Levels of Expression based on the Cell-Cycle Status” (ROLECCS) system.

We generated two different ROLECCS proteins (ROLECCS G1 and ROLECCS G2), by fusing the *Oryza sativa* TIR1 (*Os*TIR1) F-box protein, the fluorescent indicator mEmerald and the FUCCI tags Cdt1 and Geminin, respectively. The ROLECCS system exerts its targeted proteolytic activity based on a Boolean-logic computational process. In fact, the presence of the phytohormone auxin and the progression through the appropriate phase of the cell cycle are both simultaneously required to trigger the biological functions of ROLECCS proteins. As a result, the degradation of the mAID-tagged POI is temporally restricted to a specific phase of the cell cycle, and only in the presence of auxin (**Figure 1 and Figure 5**).

**Figure 5.**
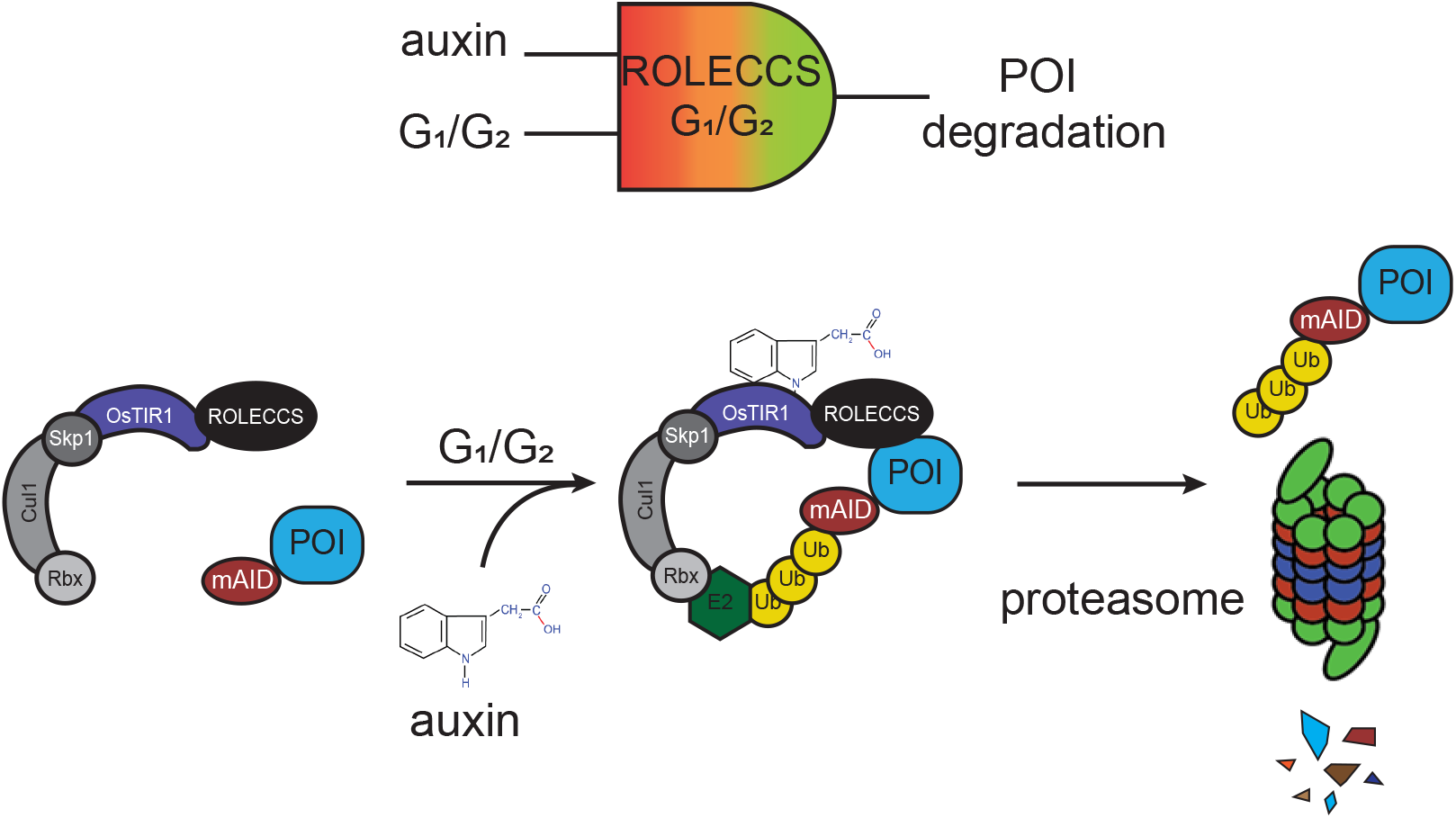
Schematic description of the ROLECCS system for cell cycle-specific targeted proteolysis. The ROLECCS system performs a Boolean logic computation. The contemporary presence of auxin and appropriate phase of the cell cycle are both simultaneously required to lead to targeted protein degradation. ROLECCS G1 and G2 are stable only through specific phases of the cell cycle (G1/early S for ROLECCS G1, late S/G2/M for ROLECCS G2), therefore their biological activity is restricted to those phases. However, auxin is required to trigger OsTIR1- mediated protein ubiquitylation, allowing proteasomal degradation of the POI only “on demand”, and only in the appropriate phase of the cell cycle.

In our first design, the ROLECCS system is specifically integrated by CRISPR/Cas9 knock-in in the *AAVS1* safe harbor genomic locus (**Figure 2**). In these settings, we observed appropriate phase-specific expression of the ROLECCS proteins. However, we noticed that exogenous targets (e.g. transiently transfected mAID-tagged POIs) were effectively down-regulated only at very short time points after the transfection. We hypothesized that this was due to molar excess of the transfected target POI in comparison with ROLECCS proteins, especially at longer time points.

For this reason, we designed All-in-One constructs, similar to others recently reported^38^, allowing the simultaneous expression of the ROLECCS proteins and their mAID-tagged targets (pLentiROLECCS), using mAID-mCherry for our tests. Our results supported the conclusion that ROLECCS proteins require to be at least in equimolar ratio to their targets to achieve consistent target degradation upon auxin treatment. Therefore, the pLentiROLECCS system is a flexible and relatively simple way to generate cell lines in which an exogenous POI can be modulated on-demand (upon auxin treatment) in specific phases of the cell cycle (**Figure 3**).

We also noticed that proteolysis of the target, although significant, was not complete, especially with the ROLECCS G2 system. One possible explanation is that, during the sorting process, cells in late G2/M phase might complete cell division and promptly enter G1, leading to re-expression of their target. Conversely, since ROLECCS G1 sorted cells are less prone to progress through cell division while in suspension, this ROLECCS protein appeared more efficient. This phenomenon could justify the efficacy differences observed between ROLECCS G1 and ROLECCS G2 (**Figure 3G-H**).

Another possibility is that, to maximize the efficacy of ROLECCS variants, ROLECCS proteins should be further modified to enhance their localization in the same cellular compartment as their target. In fact, we observed that ROLECCS G1 and ROLECCS G2 markedly localized to the nucleus, similarly to the FUCCI tags^19^. However, we also detected noticeable levels of ROLECCS proteins in the cytoplasm. This phenomenon could be at least in part explained by the ubiquitous localization of the Skp1/Cul1/Rbx/OsTIR1 complex, which could counteract the nuclear localization stimulated by the FUCCI tags. In fact, as shown in **Figure 2C**, in the absence of Cdt1 or GEM tags, OsTIR1-Emerald mainly localized in the cytoplasm. Conversely, addition of the FUCCI tags led to a marked, although not complete, nuclear localization. It is therefore tempting to speculate that the addition of Nuclear Localization or Export Signals (NLS and NES) to the ROLECCS systems could allow an unprecedented spatio-temporal control of protein levels. The use of pLentiROLECCS may lead to the expression of the POI at levels that are several folds higher than the endogenous protein, because of the exogenous strong promoter (CMV) controlling the expression of the ROLECCS-P2A-POI fusion. However, our ultimate goal was to generate a system to synthetically control endogenous protein levels based only on the cell cycle status, minimizing potential artificial factors such as the use of an exogenous promoter.

Therefore, as proof-of-principle, we attempted to regulate the levels of a protein encoded by an endogenous gene, fused by CRISPR/Cas9 knock-in with the mAID tag. We chose the gene *TP53* because of its well-known role in the regulation of cell cycle progression^42^. Moreover, gene editing of this gene with the mAID tag was previously reported^33^. However, we hypothesized that, using the ROLECCS system, we could deplete TP53 protein levels in specific phases of the cell cycle. Here, we show that cell cycle-specific TP53 degradation was effectively achieved with both the ROLECCS G1 and the ROLECCS G2 systems (**Figure 4**). The molecular and biological implications of the selective degradation of TP53 in dividing or not dividing cells are currently being explored.

The ROLECCS system described here is the prototype of multiple potential cell cycle phase-specific degron technologies. For example, the choice of different FUCCI tags, as described previously, could lead to a more accurate phase-specific protein degradation. In fact, the FUCCI tags used for the ROLECCS proteins in the present study are partially co-expressed in late- G1/early-S phase. However, it has been previously demonstrated the different Cdt1 domains could be used to achieve a sharp down-regulation of the ROLECCS system at the G1/S transition^24^.

A major weakness for the OsTIR1-based mAIDs is their potential leakiness, which could lead to protein downregulation in the absence of auxin^31, 45^. One potential solution is represented by the use of other F-box proteins, such as the *Arabidopsis Thaliana* AFB2 protein, which demonstrated minimal basal depletion of target proteins in the absence of auxin^45^. Alternatively, negative regulators of the AID system, such as the auxin response transcription factor ARF, can be leveraged to limit chronic target protein depletion in the absence of auxin^49^. More recently, a point mutant of OsTIR1 (F74G) was reported, establishing the mAID version 2 (mAID2) system, which does not respond to natural auxin but only to a synthetic ligand (5-Ph-IAA). Interestingly, this point mutant displayed no detectable leaky degradation of the target, was responsive to 670-times lower concentration of the ligand^31^ and it was functional also *in vivo* using mouse models.

Protein biological functions and cell cycle progression are intimately connected and reciprocally affected. Hence, the cell cycle status should be taken into account for the study of any biological phenomenon. Thanks to its phase specificity, rapidity, reversibility, and low overall perturbation of other biological processes, the ROLECCS technology represents a unique tool for the investigation of biological phenomena and their relationship with the cell cycle progression.

## Methods

### Plasmids generation

All the plasmids used in this study were generated by Gibson Assembly using NEBuilder® HiFi DNA Assembly Master Mix (New England Biolabs, E2621L) as per manufactory’s instructions. To construct pAAVS1-ROLECCS AS, pAAVS1-ROLECCS G1, and pAAVS1-ROLECCS G2 plasmids, multiple fragments were PCR amplified from different donor plasmids and assembled as follow: pMK232 CMV-OsTIR1-PURO (Addgene #72834^33^) was used as donor plasmid for the expression of OsTIR1 from the *AAVS1* locus, the mEmerald tag was PCR amplified from mEmerald-PLK1-N-16 vector (Addgene #54234; http://n2t.net/addgene:54234 ; RRID:Addgene_54234), while pEN435 - pCAGGS-TagBFP-hGeminin-2A-mCherry-hCdt1- rbgpA-Frt-PGK-EM7-PuroR-bpA-Frt Tigre targeting (Addgene #92139^25^) was used as template for both hGeminin and hCdt1 tags. The vector for the mAID-mCherry expression was generated using the pEGFP-C1 backbone (Clontech), replacing the GFP gene with the mAID-mCherry cassette derived from pMK292 mAID-mCherry2-NeoR (Addgene #72830^33^). The bicistronic lentiviral vectors for ROLECCS AS, ROLECCS G1, ROLECCS G2, and mAID-mCherry expression were similarly obtained, although linker sequences (P2A) were synthesized (IDT) and cloned by Gibson assembly.

To generate the donor plasmids for *TP53* editing, the genomic region (∼2000bp) encoding for the natural stop codon of *TP53* was first cloned into the pUC19 vector (New England Biolabs, N3041S) by Gibson assembly. More specifically, genomic DNA from H460 cell line (TP53 wild-type)^50^ was used as template to amplify the *TP53* genomic region (Chromosome 17: 7,668,421- 7,687,490, Transcript: TP53-201 ENST00000269305.9) of 1 kb upstream and 1 kb downstream the *TP53* translation stop codon. These regions were further used as homology arms for HDR-mediated CRISPR/Cas9-mediated knock-ins. Secondly, the homology arms containing plasmid was mutated using QuikChange II XL Site-Directed Mutagenesis Kit (Agilent, #200522) to delete the single-guide RNA (sgRNA) recognition sequence, to prevent Cas9 from re-cutting after homology-directed repair-mediated insertion at the desired genetic locus. Finally the mAID-mCherry cassette containing a selection marker was amplified from pMK292 mAID-mCherry2- NeoR (Addgene #72830^33^) or pMK293 mAID-mCherry2-Hygro (Addgene #72831^33^) and inserted between the homology arms (about 1000bp each), replacing the *TP53* stop codon, making sure that the tags sequences were cloned in frame with the gene of interest, in order to generate a fusion protein. A schematic overview of the donor vectors is presented in **Figure 4A**.

To construct the CRISPR/Cas9 *TP53* gene targeting vector, a single-guide RNA (sgRNA) (5’- ACTGACAGCCTCCCACCCCC-3’) was designed (http://crispr.mit.edu) to specifically target TP53 translation stop site and it was cloned into pX330-U6-Chimeric_BB-CBh-hSpCas9-hGem (1/110) (Addgene #71707) according to the protocol of Ran et al.^51^. The same protocol was followed to clone the sgRNA used for the ROLECCS transgene insertion in the *AAVS1* locus^33^ (PMID: 27052166), into the pX330-U6-Chimeric_BB-CBh-hSpCas9-hGem (1/110)^52^.

All the plasmids will be deposited on Addgene or are available from the investigators upon kind request.

### Cell Culture, Transfection, and Clones Isolation

The HCT116 and HEK-293T (HEK-293) cell lines were purchased from American Type Culture Collection (ATCC CCL-247; ATCC CRL-11268) and were cultured in RPMI-1640 medium (Millipore Sigma) supplemented with 10% FBS (Millipore Sigma). Cells were grown in a 37 °C humid incubator with 5% CO^2^. Identity of cell lines was validated upon and after the establishment of stable clones by STR profiling. To generate HEK-293 constitutively expressing pAAVS1- ROLECCS AS, pAAVS1-ROLECCS G1, and pAAVS1-ROLECCS G2, 2×10^5^ cells were plated in a six-well plate, and 24h later CRISPR/Cas9 and donor plasmids were transfected using Lipofectamine™ 2000 Transfection Reagent (Thermo Fisher Scientific, #11668019) in Opti- MEM™ I Reduced Serum Medium (Thermo Fisher Scientific, #31985070). Cells were then grown up to a subconfluent T175 cm^2^ flask in medium supplemented with 10% FBS and, about ten days after transfection, antibiotic selection was started using 2 µg/ml Puromycin Dihydrochloride (Thermo Fisher Scientific, A1113803) in medium. Resistant populations were expanded up to a subconfluent T175 cm^2^ flask and cells were sorted and collected at FACSAria II (Becton Dickinson) for brightest mEmerald expression, using 488nm laser excitation. Parental HEK-293 cells were used as negative control of mEmerald expression to design the gates. Single cell clones were grown in 96-well plates and screened by WB for ROLECCS AS, G1 and G2 expression. Two different clones for each constructs were chosen among the ones with comparable ROLECCS expression. To obtain HEK-293 constitutively expressing pLENTIROLECCS AS, G1 or G2, 2×10^5^ cells were plated in a six-well plate and the next day they were transfected with plasmids using Lipofectamine™ 2000 Transfection Reagent (in Opti-MEM™ I Reduced Serum Medium). After 48 hours from transfection, transfected cells selection was started using 5 µg/ml Blasticidin S HCl (Thermo Fisher Scientific, A1113903) in complete medium. Resistant population was expanded up to a subconfluent T175 cm^2^ flask and double positive cells for mEmerald and mCherry expression were sorted and collected at FACSAria II, using 488nm and 561 nm laser excitations. Parental HEK-293 cells were used as negative control of mEmerald and mCherry expression to design the gates. Single cell clones were grown in 96 well plates and screened by WB for ROLECCS AS, G1 or G2 and mAID-mCherry expression. Two different clones for each constructs were chosen among the ones with comparable proteins expression.

To generate *TP53-*edited HCT116 cells, 2×10^5^ cells were plated in a six-well plate and CRISPR/Cas9 along with donor plasmids were transfected using Lipofectamine™ 3000 Transfection Reagent in Opti-MEM™ I Reduced Serum Medium. Cells were then grown up to a subconfluent T175 cm^2^ flask in medium supplemented with 10% FBS and, about ten days after transfection, antibiotic selection was started using 700 µg/ml G418 Sulfate (Thermo Fisher Scientific, #10131035) and 100 µg/ml Hygromycin B (Thermo Fisher Scientific, 10687010) in medium. Double resistant population was expanded up to a full T175 cm^2^ flask and cells were sorted and collected at FACSAria II for brightest mCherry expression, using 561 nm laser excitation. Parental HCT116 cells were used as negative control of mCherry expression to design the gates. Single cell clones were grown in 96-well plates and screened by WB for TP53-mAID-mCherry expression. Two different clones were chosen among the ones with homozygous expression of TP53 edited protein. After genotyping (see next section), the two clones were gene edited to constitutively express the pAAVS1-ROLECCS variants, following the transfection and selection protocol aforementioned for HEK-293 cells.

To test biological functionality of ROLECCS constructs, HEK-293 *AAVS1*-integrated clones were transfected with 150 ng of mAID-mCherry expressing plasmid (see “Plasmid Generation” section) using Lipofectamine 2000. After 7 hours from transfection, cells were treated with 500 µM auxin for one hour. Cells were then collected and processed for WB analysis.

### Genomic PCR and genotyping

To obtain genomic DNA, cell pellets were resuspended in lysis solution (100 mM Tris-HCl [pH 8.0], 200 mM NaCl, 5 mM EDTA, 1% SDS, and 0.6 mg/ml proteinase K), and incubated at 55°C overnight. After ethanol and sodium acetate precipitation, DNA pellets were washed in 70% ethanol and resuspended in water. The DNA solution was incubated at 60 °C for 15 min and at least 1 hour at room temperature before proceeding. Genomic PCR was performed using Q5® High-Fidelity 2X Master Mix (New England Biolabs, M0492L) according to the manufacturer’s instruction. To genotype *TP53-*edited HCT116 clones, purified DNA (50ng) was analyzed by PCR to verify biallelic insertion of mAID-mCherry tag along with antibiotic resistance, in the right genetic locus, using the following primers: A (5’-TTGGAACTCAAGGATGCCCAGG-3’); B (5’- CATGGCCAGCCAACTTTTGCAT-3’); E (5’ATTTCGGCTCCAACAATGTC-3’): F (5’- TGCTCCTGCCGAGAAAGTAT-3’); C (5’-GGATGTTCCGAGAGCTGAAT-3’); D (5’- GAAGAACGTGATGGTTTC-3’) (**Supplementary Figure 5**). PCR products were loaded on 1 or 2% agarose TBE with ethidium bromide gel, along with 100bp or 1kb DNA ladders (Thermo Fischer Scientifc) to verify correct amplicons length and then purified using QIAquick PCR Purification Kit (Qiagen). Purified PCR DNA was submitted for sequencing to assess the frame and the integrity of the edited sequence, using the following primers: pSHALfwd (5’- ACTGAATACAGCCAGA-3’); pSHALrev (5’-ACTGAATACAGCCAGA-3’); A (5’- TTGGAACTCAAGGATGCCCAGG-3’); mAIDfwd (5’-GAAGAACGTGATGGTTTC-3); D (5’-GAAGAACGTGATGGTTTC-3’).

### Treatments

To induce the degradation of mAID-fused proteins, cells were treated with 500 µM indole-3-acetic acid (auxin, IAA, dissolved 500 mM in water) (Millipore Sigma, I5148) for 1 hour, unless otherwise stated. To suppress the partial degradation of TP53-mAID-mCherry in HCTT116 constitutively expressing pAAVS1-ROLECCS AS, pAAVS1-ROLECCS G1 and pAAVS1- ROLECCS G2 control cells were pretreated overnight with 200 µM auxinole (BioAcademia, Inc., Japan; #30–001, dissolved 200 mM in DMSO) and maintained in auxinole for the duration of the experiments where indicated.

To induce TP53 activation, cells were treated with 20 µM cis-Diamineplatinum(II) dichloride (cisplatin, stock diluted in 0.9% NaCl, Millipore Sigma, #479306) for 48 hours.

### Protein extractions, subcellular fractionations, and western blots

Total protein extractions were performed as follows: cells were collected and washed with PBS before adding adequate amount of lysis buffer (1% NP-40, 1mM EDTA pH 8.00, 50Mm Tris-HCl pH 7.5, 150 mM NaCl) containing a protease and phosphatase inhibitor cocktail (cOmplete™, EDTA-free Protease Inhibitor Cocktail; PhosSTOP™ inhibitor tablets, Millipore Sigma). Subcellular protein fractionations were performed using the NE-PER™ Nuclear and Cytoplasmic Extraction Reagents (Thermo Fisher Scientific, 78833), according to manufacturer’s instructions. Protein concentration was checked by Bradford assay (Biorad). After denaturation at 100°C for 5 min, equal amounts of proteins (2.5-25 µg) were separated using SDS-PAGE, loading samples on TGX™ Precast Protein Gels (Bio-Rad). Proteins were transferred to 0.45 µm nitrocellulose membrane (Biorad) and blocked in 5% non-fat milk or BSA/TBST for 1 hour at room temperature. Membranes were probed with primary antibodies at 4°C overnight and subsequently incubated with a secondary antibody at room temperature for 1 hour. Antibodies used were anti-GFP (B-2) (Santa Cruz Biotechnology, sc-9996), anti-TP53 (DO-1) (Santa Cruz Biotechnology, sc-126), anti-GAPDH (14C10) Rabbit mAb (HRP Conjugate) (Cell Signaling Technology, #3683), anti-Cyclin B1 (D5C10) XP Rabbit mAb, (Cell Signaling Technology, #12231), anti-CDT1 (D10F11) Rabbit, (Cell Signaling Technology, #8064), anti- p21 Waf1/Cip1 (12D1) Rabbit mAb, (Cell Signaling Technology, #2947), anti-mCherry (Millipore Sigma, AB356482), anti-nucleophosmin (Cell Signaling Technology, #3542), anti-vinculin monoclonal Antibody (VLN01, Thermo Fisher # MA5-11690). Secondary antibodies were HRP-conjugated anti-mouse or rabbit IgG from Ge (LI-COR Bioscience). Detection was performed using Immobilon Forte Western HRP substrate (Millipore Sigma) or the SuperSignal™ West Femto Maximum Sensitivity Substrate (Thermo Fisher Scientific) and X-ray blue films, in dark room. Blots in the same figure/panel were probed multiple times with the indicated antibodies and, when necessary, membranes were stripped using Restore™ PLUS Western Blot Stripping Buffer (Thermo Fisher Scientific, 46428) and re-probed with the desired antibody. Images were acquired using standard protocols and Western Blot bands were quantified using ImageJ software (U. S. National Institutes of Health, Bethesda, Maryland, USA).

### Gating and sorting strategy

For FACS sorting, all samples were washed once in 1X PBS and resuspended in sorting buffer (1X PBS, 1mM EDTA, 24mM HEPES pH 7.0, 1% FETAL Bovine Serum (Heat-Inactivated), 0.2 µm filtered) before performing flow cytometric analysis using FACSAria II (Becton Dickinson). After gating cells to exclude debris, dead cells, and doublets, cells were plotted for SSC-A and FITC-A (mEmerald expression). Three different gates, labelled GFP^low^, GFP^med^, and GFP^high^, were designed to identify three distinct cell populations with different intensity of FITC (that is mEmerald expression). We also took into consideration the SSC as indicator of cells complexity to help recognize cells progressing into different phases of the cell cycle, especially to discriminate cells in G2 phase. Therefore, in the instance of sorting ROLECCS G1 cells, the GFP^high^ gate include cells with high FITC intensity and low SSC, as these are cells expected to be in G1 phase and so with relatively low cellular complexity and high expression of OsTIR1-mEmerald-Cdt1 protein. In the case of ROLECCS G2 cells, the GFP^high^ population is the one with high SSC and high FITC level as the OsTIR1-mEmerald-Geminin expression increases in S/G2 as the cellular complexity does. GFP^low^ and GFP^med^ gates are here being indicated and analyzed as counter-proof that our reporters are not expressed in the not specific cell cycle phase, therefore, when cells are sorted for low or medium expression of mEmerald (FITC) they are not in the cell cycle phase under study. Sorting cells expressing ROLECCS G1 or ROLECCS G2 is intended as sorting for GFP^high^ cell populations, as the portion of the asynchronously growing cells, where solely happens the on demand degradation of the POI. Representative gating strategies are shown in **Supplementary Figures 2** and **6**.

During sorting experiments with HEK-293 expressing pLentiROLECCS, red fluorescent intensity from mAID-mCherry was simultaneously acquired on GFP^low^, GFP^med^, and GFP^high^ populations, using 561 nm laser excitation. Flow cytometry data were analyzed using FlowJo™ Software_v 10.6.1. (Ashland, OR: Becton, Dickinson and Company; 2019) analysis software and presented as PE-Texas Red-A intensity *versus* counts.

### Cell cycle analysis using propiudium iodide DNA staining

Cells were washed in 1X PBS (Millipore Sigma) before being fixed in 70% ethanol at -20 °C for at least 2 hours. After fixation, cells were pelleted and washed once in 1X PBS and then resuspended in staining solution (PBS containing 10 mg/mL propidium iodide (Thermo Fisher Scientific), 0.05% Triton X-100 (Millipore), 2.5 µg/mL RNAse A (Thermo Fisher Scientific)). After samples incubation at 37 °C for 30 min protected from light, flow cytometry analysis was performed using FACSCalibur Flow Cytometer (Becton Dickinson), modelling at least 5000 events per sample. Results were analyzed using ModFit software, v5.0 (Verity Software House).

### Cell imaging

For live cell imaging experiments, HEK-293 ROLECCS *AAVS1*-integrated clones were plated in 96-well plates and time-lapse analyses were performed using IncuCyte® S3 Live-Cell Analysis System (Essen BioScience). Images were acquired every 15 minutes using bright-field and GFP channels. Data were exported and presented as unprocessed.

Images of fluorescent features of HEK-293 ROLECCS *AAVS*- integrated clones were obtained at Zeiss Axioskop 40 Microscope, using Zen Pro software (ZEISS). Cells were plated in cellview cell culture dish (glass bottom, Greiner Bio-One, 627870) and nuclear counterstaing was performed adding one drop of NucBlue™ Live ReadyProbes™ Reagent (Invitrogen, R37065) to the medium. DAPI (blue) and mEmerald (green) were acquired using EGFP and DAPI channels at 63x magnification.

### Statistical analysis and data availability

All the experiments are representative of at least two independent experiments (technical and/or biological replicates). The number of replicates for each experiment is specified in the relative figure legend. For statistical analysis, two-tailed t-test was performed and data were considered statistically significant for *p<*0.05.

Original unprocessed data used for the preparation of the manuscript are available upon kind request and will be uploaded on Mendeley Data upon paper submission.

The full list of used primers and reagents used in this study, as well as the plasmids generated is available as supplementary files.

## Acknowledgements

We thank Dr. Emanuele Cocucci for the scientific discussions during the preparation of this manuscript. We also thank the countless investigators who made their plasmids and reagents available through public repositories such as Addgene. This work was supported by seed funds from The Ohio State University Comprehensive Cancer Center (DP, CMC), Pelotonia (DP) and NCI-NIH (NIH R35CA197706) (CMC). The Gene Editing Shared Resource, the Flow Cytometry Shared Resource and the Genomics Shared Resource that contributed to this study are supported by the Cancer Center Support Grant P30CA016058.

## Author Contributions

M.C., D.P. and C.M.C. conceived and designed the project. M.C., D.P., A.T., W.O.M. conceived and planned the experiments. M.C., D.P., A.T., J.M., G.L.R.V., C.L., W.O.M. executed the experiments. M.C., D.P., A.T., G.L.R.V., C.L., W.O.M. analyzed the data. B.M. provided technical support for flow cytometry experiments and cell sorting. M.C. and D.P., wrote the manuscript. A.T., J.M., G.L.R.V., C.L., W.O.M., V.C. and C.M.C. provided critical feedback and contributed to the final version of the manuscript.

---

**Supplementary Figure 1.**
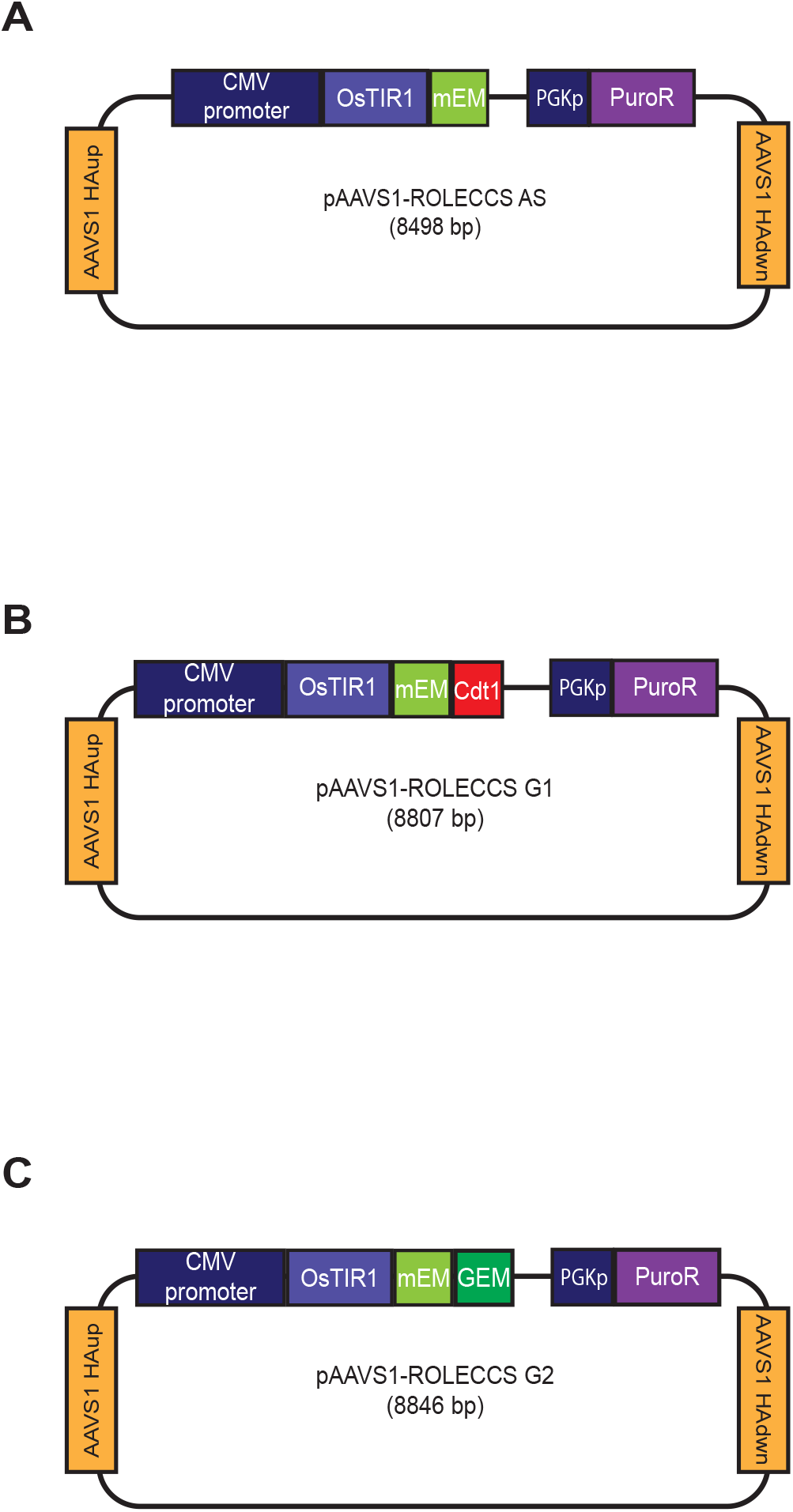
ROLECCS plasmids for *AAVS1* targeted insertion. (A-C) Schematic depiction of pAAVS1-ROLECCS vectors for *AAVS1*-targeted insertion. The ROLECCS cassette (OsTR1-mEmerald-/Cdt1/Geminin), under the CMV promoter is cloned between the two AVVS1 Homology Arms (AAVS1 HA) up and down (dwn), along with the gene for puromycin resistance, controlled by the human PGK promoter.

**Supplementary Figure 2.**
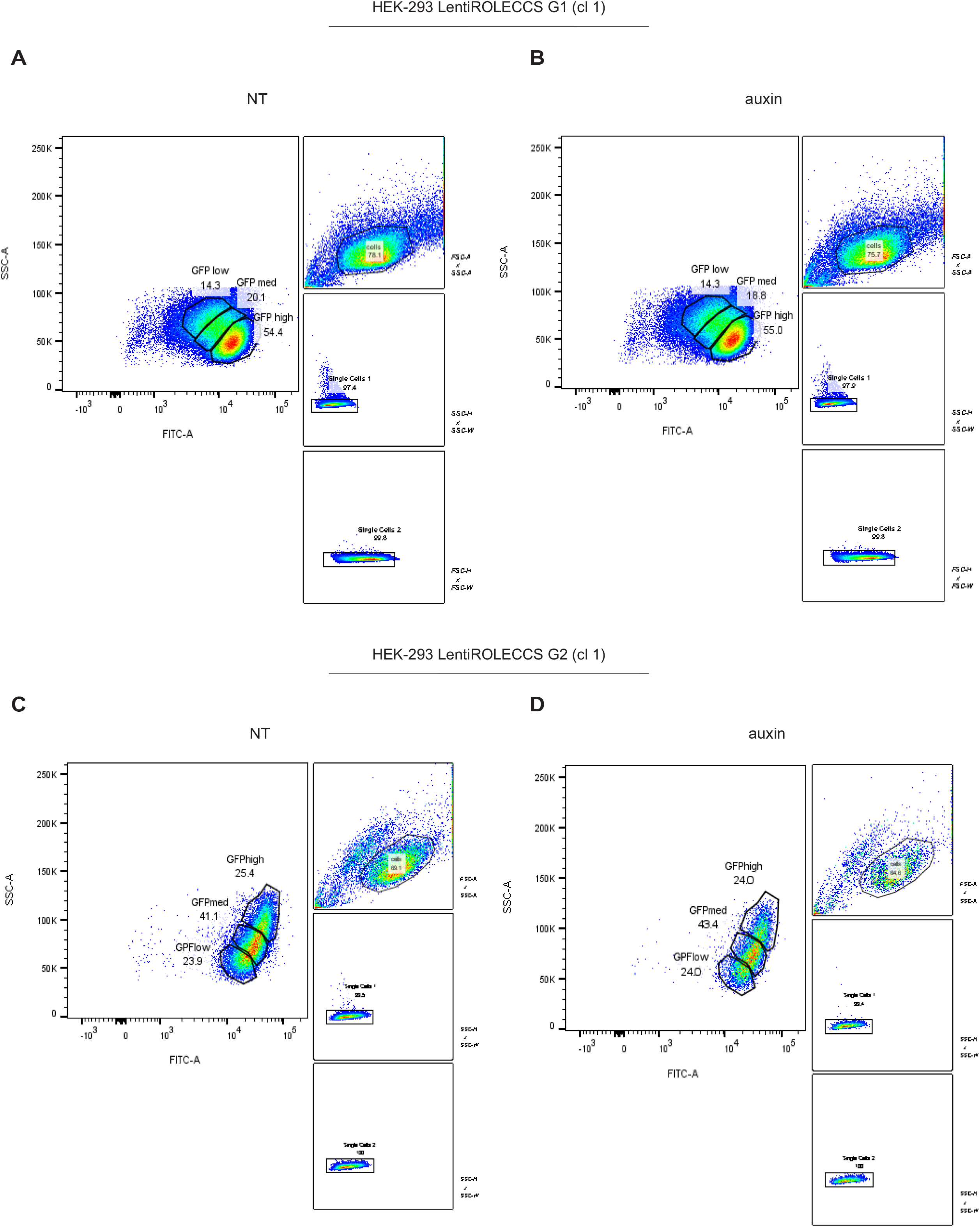
Gating strategy for HEK-293 LentiROLECSS G1 and G2 sorting. (A-D) Cells were plotted for side scatter (SSC-A) and FITC-A (mEmerald intensity), after gating to exclude debris, dead cells, and doublets, as shown by ancestry plots reported on left side of SSC-A *versus* FITC-A panel. Three different gates, labelled GFP^low^, GFP^med^, and GFP^high^, were designed to identify three distinct cell populations with different intensity of FITC, taking into consideration the SSC as indicator of cells complexity to help recognize cells progressing into different phases of the cell cycle. Not treated (NT) and auxin treated cells were sorted with the same strategy.

**Supplementary Figure 3.**
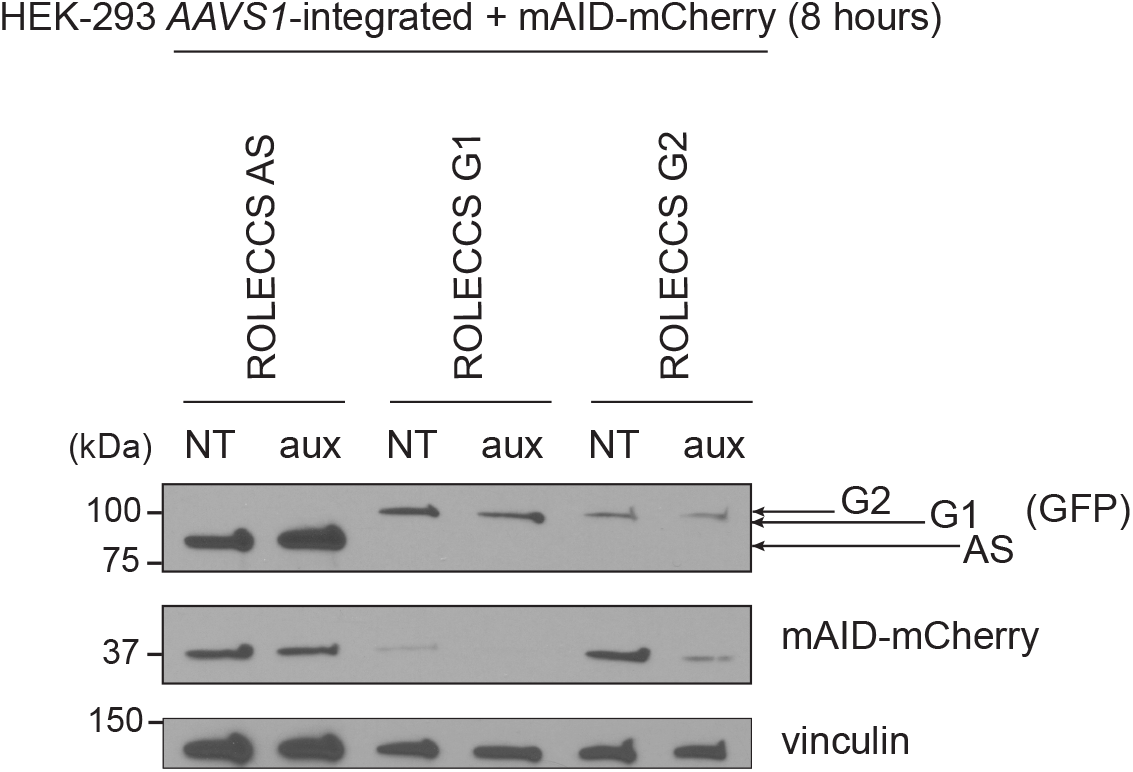
Biological activity of ROLECCS proteins on transiently transfected target. WB analysis of mAID-mCherry degradation upon auxin treatment. HEK-293 *AAVS1*-integrated clones, expressing ROLECCS AS/G1/G2, were transfected with mAID-mCherry expressing plasmid. After 7 hours from transfection, cells were treated with 500 µM auxin (aux) or not treated (NT) for 1 hour (total 8h), without removing the transfection mix. Anti-GFP antibody was used to detect ROLECCS variants, anti-mCherry antibody was used to detect mAID-mCherry and vinculin was used as loading control.

**Supplementary Figure 4.**
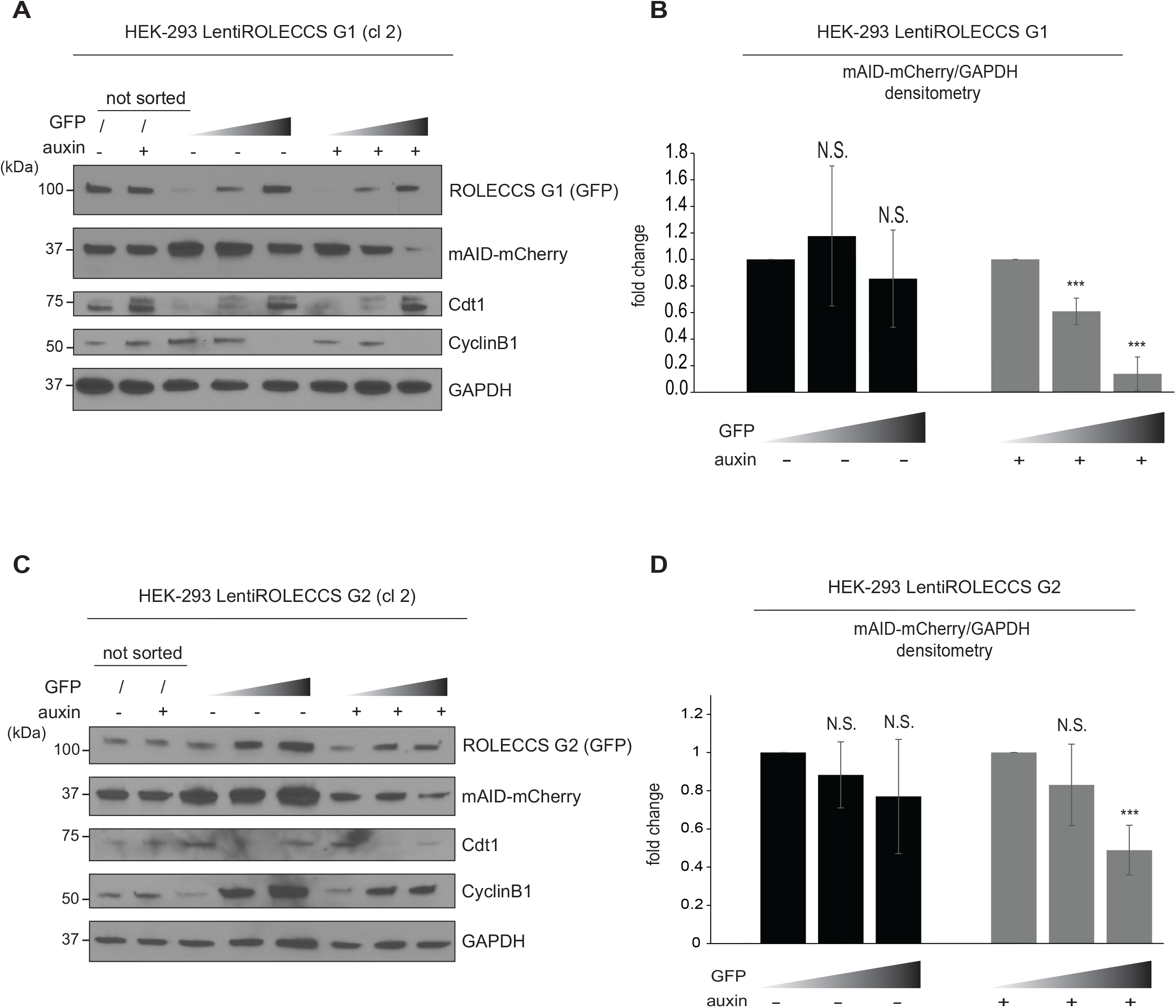
Supporting data of functionality of ROLECCS G1 and G2 proteins. (A-B) WB analysis (A) and densitometric quantification (B) of HEK-293 cells transfected with pLentiROLECCS G1, clone 2 (cl 2) after sorting. Cells were treated with auxin or left untreated for one hour and then sorted for increasing GFP intensity (ascending grey gradient triangle). Membrane was probed with anti-GFP antibody for ROLECCS G1 detection and mCherry antibody for mAID-mCherry detection. Cdt1 and CyclinB1 were used as G1 phase and G2 phase-specific markers, respectively. GAPDH was used as loading control. Not sorted (unsorted) cells were loaded for comparison. Densitometric quantification of mAID-mCherry was normalized on GAPDH intensity and relative quantification *versus* GFP^low^ sorted population is reported. (C-D) WB analysis (C) and densitometric quantification of HEK-293 cells transfected with pLentiROLECCS G2, clone 2 (cl 2) after sorting. Treatments, sortings and antibodies are the same as (A). Densitometric quantification was performed as in (B). Blots are representative of 2 independent experiments. Densitometric analyses are the average of 2 independent experiments on 2 different clones (n=4).

**Supplementary Figure 5.**
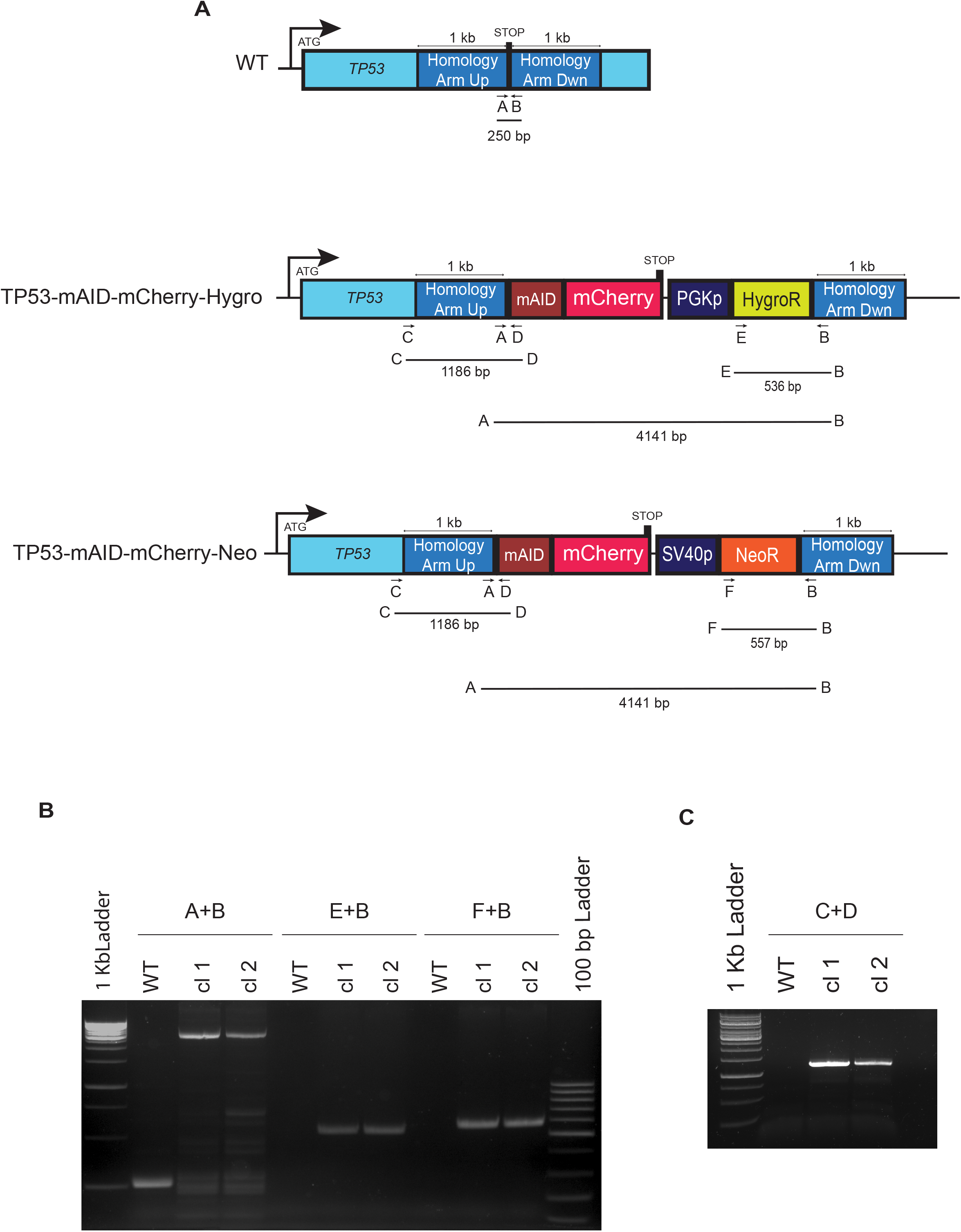
Homozygous editing of the *TP53* locus in HCT116 cells. (A) Schematic illustration of wild type (WT) and edited (*TP*53-mAID-mCherry-Hygro/Neo) alleles. Primers for genotyping are indicated by arrows and capital letters and reported under their annealing site. Expected PCR products are specified by black bars, with corresponding amplicon length. (B) HCT116 clone 1 and 2 (cl 1 and 2) have inserted mAID-mCherry fusion cassette through homologous recombination with Homology Arms, as shown by PCR with primers set A+B. The insertion of the cassette is on both alleles as shown by PCR products with primers sets E+B (for HygroR bearing Donor plasmid) and F+B (for NeoR bearing Donor plasmid). (C) Genomic PCR to verify that the insertion is at the right *TP*53 genomic *locus* (primer set A+D). HCT116 wild type cells (WT) were used as control and 1 Kb and 100 bp ladders were loaded along with PCR products on 1% gel.

**Supplementary Figure 6.**
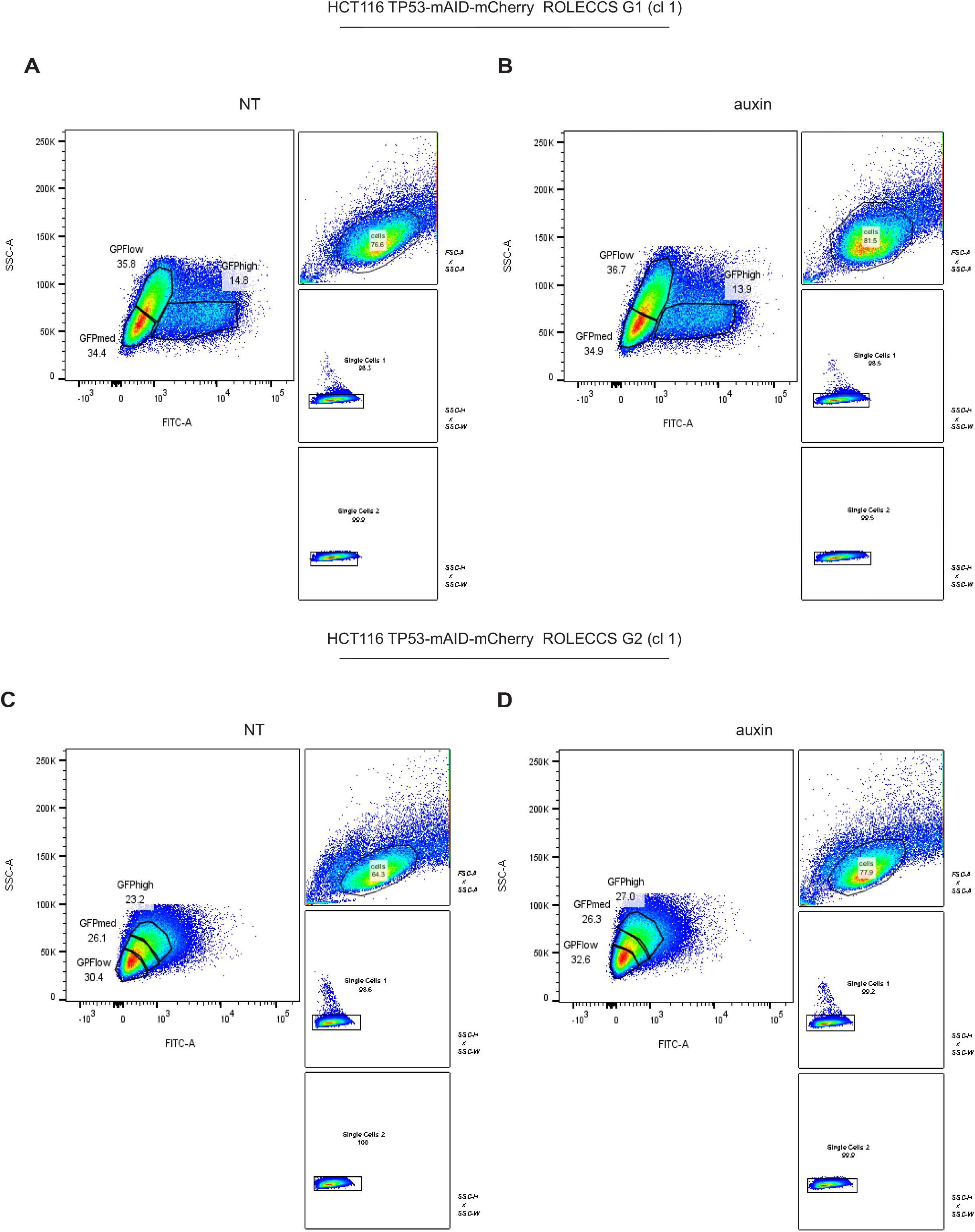
Gating strategy for HCT116 TP53-mAID-mCherry ROLECSS G1 and G2 sorting. (A-D) Cells were plotted for side scatter (SSC-A) and FITC-A (mEmerald intensity), after gating to exclude debris, dead cells, and doublets, as shown by ancestry plots reported on left side of SSC-A *versus* FITC-A panel. Three different gates, labelled GFP^low^, GFP^med^ and GFP^high^, were designed to identify three distinct cell populations with different intensity of FITC, taking into consideration the SSC as indicator of cells complexity to help recognize cells progressing into different phases of the cell cycle. Not treated (NT) or auxin treated cells were sorted using the same strategy.

**Supplementary Figure 7.**
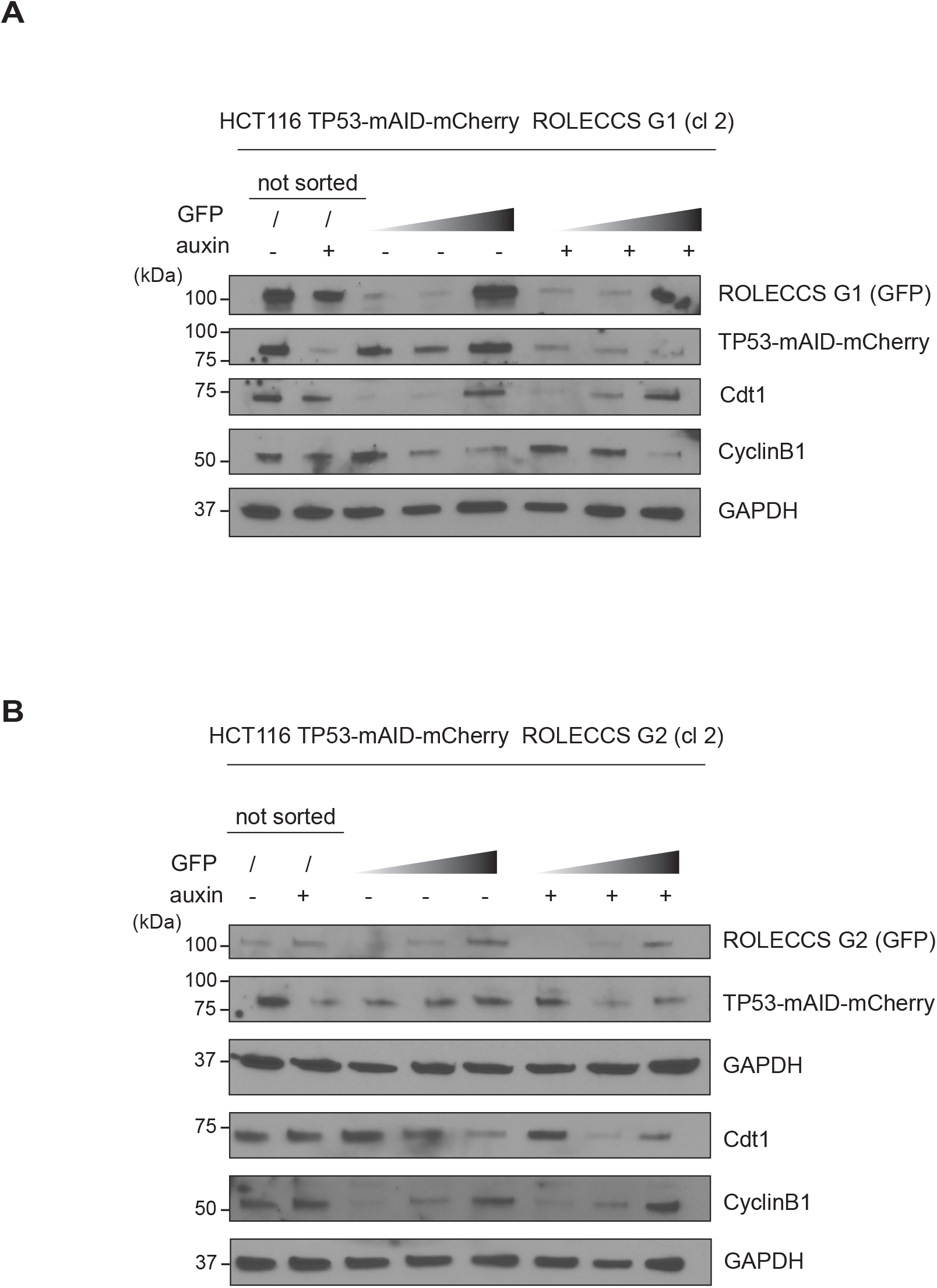
Supporting data of functionality of ROLECCS system on endogenous proteins. (A) WB analysis of HCT116 TP53-mAID-mCherry *AAVS1*-edited with ROLECCS G1, clone 2 (cl 2) after sorting. Cells were treated with auxin or left untreated for one hour and then sorted for GFP intensity (ascending grey gradient triangle). Membrane was probed with anti-GFP antibody for ROLECCS G1 detection and mCherry antibody for TP53-mAID-mCherry (TP53-mAID-mCh) detection. Cdt1 and CyclinB1 were used as G1 phase and G2 phase specific markers, respectively. GAPDH was used as loading control. Not sorted (unsorted) cells were loaded for comparison. (B) WB analysis of HCT116 TP53-mAID-mCherry *AAVS1*-edited with ROLECCS G2, clone 2 (cl 2) after sorting. Treatments, sortings, and antibodies are the same as in panel A. Blots are representative of two independent experiments.

